# EZHIP boosts neuronal-like synaptic gene programs and depresses polyamine metabolism

**DOI:** 10.1101/2025.05.08.652913

**Authors:** Elham Hasheminasabgorji, Huey-Miin Chen, Taylor A Gatesman, Subhi Talal Younes, Gabrielle A Nobles, Farhang Jaryani, Heather Mao, Kwanha Yu, Benjamin Deneen, Wee Yong, Michael D Taylor, Sameer Agnihotri, Marco Gallo

## Abstract

It is currently understood that the characteristic loss of the repressive histone mark H3K27me3 in PFA ependymoma and diffuse midline glioma (DMG) are caused by complementary mechanisms mediated by EZHIP and the oncohistone H3K27M, respectively. To support the complementarity of these mechanisms, rare H3K27M-negative DMGs express EZHIP. Interestingly, EZHIP is one of the few genes recurrently mutated in PFA. The significance of EZHIP mutations in PFA, and whether EZHIP has wider functions in addition to repression of H3K27me3 deposition, are not known. Here, we investigated the mutational landscape of EZHIP in pediatric brain tumors. We found that EZHIP mutations occur not only in PFA, but also in rare medulloblastoma and pediatric high-grade glioma (HGG), including in H3K27-positive DMG. Contrary to current expectations, we show that mutant EZHIP is expressed in H3K27M-positive DMG. All the EZHIP-mutated HGG cases also have EGFR mutations. Further, we pursued better understanding of the function of EZHIP by expressing it in human-derived neural models. Our transcriptomic analyses indicate that EZHIP expression potentiates neuronal-like gene programs associated with synaptic function. Metabolomics data indicate that EZHIP leads to repression of methionine and polyamine metabolism, suggesting links between metabolic and epigenetic changes that are observed in PFA. Collectively, our results expand the repertoire of tumor types known to harbor EZHIP mutations and shed light on EZHIP-dependent metabolic and transcriptional programs in relevant neural models.

## INTRODUCTION

The function of EZHIP (enhancer of zeste inhibitory protein) is not yet clearly and completely defined in developmental and malignant cellular contexts. *EZHIP* is located on chromosome X in a genomic region with high frequency of repeat sequences. It is only present in some placental mammals^1,2^ and its evolutionary origin is still unresolved. Its expression starts in primordial germ cells following their migration to the genital ridge at 7 weeks post-conception and it then persists in testes and ovarian follicles^3^.

EZHIP has no discernible conserved protein domain and its structure is predicted to be mostly disordered. Consequently, it has been difficult to infer and test potential hypothesis-based functions for this protein. EZHIP was initially identified in human and mouse models as an uncharacterized protein (previously known as CXorf67) interacting with Polycomb Repressive Complex 2 (PRC2)^1,2,4^, which deposits the repressive histone mark H3K27me3. This finding suggested a potential function for EZHIP in regulating the function of PRC2. In agreement, knockout mice have increased levels of the repressive histone mark H3K27me3, but no other overt phenotypes except for decreased fertility in females^1^. This finding is consistent with biochemical studies in human cell lines that found that EZHIP interacts with PRC2 and inhibits its H3K27 methyl-transferase activity^2,5^. Further biochemical characterization provided evidence that EZHIP blocks the interaction between PALB2 and BRCA2 and consequently inhibits homologous recombination-mediated DNA repair in cancer cell lines^6^. Beside for its involvement in regulating H3K27me3 deposition and in DNA repair, no other function has been identified for EZHIP.

There has been a considerable amount of interest in EZHIP because it is expressed in posterior fossa type A (PFA) ependymoma^6^ and few other malignancies^7^. PFA is a malignant tumor that arises in the infratentorial region of the brain. It is mostly diagnosed in infants and very young children, with an average age at diagnosis of 2.5 years^8^. Incidence of this tumor type is skewed toward males, with a male to female ratio of 1.6. Two thirds of PFA patients survive at 5 years and only half of patients survive at 10 years with current treatment approaches, which consist of maximal tumor resection and radiation. No chemotherapy has so far shown efficacy in over three decades of clinical trials^8^. Unfortunately, radiation treatment may lead to severe cognitive and developmental sequelae in long-term survivors, highlighting the need for precise therapies that exploit molecular mechanisms that are essential for PFA cellular fitness.

96% of PFA tumors express EZHIP^9^, which represents a true hallmark for this cancer type. We have recently shown that EZHIP expression is associated with global restructuring of the 3D genome of PFA cells and results in the emergence of TULIPs (Type B Ultra Long-range Interactions in PFA), which are regions of extreme chromatin compaction that interact with each other with high strength^10^. In addition to specific 3D genome configurations, PFA tumors also present with a CpG island methylator phenotype (CIMP), which consists of high methylation levels at a subset of CpG islands across the genome^11^. It is possible that the CIMP phenotype of PFA cells reflects specific metabolic states in this tumor type^12^, given that DNA methylation is influenced by metabolic dynamics.

Although PFA has characteristic DNA methylation profiles and 3D genome architecture, its linear genome is relatively stable, with only chromosome 1q gains and 6q losses occurring with appreciable frequency, and overall low mutation burden and rare coding mutations^11^. *EZHIP* is the most recurrently mutated gene in PFA, with about 10% of patients carrying mutations in this gene^13^. The functional significance of these mutations is not currently known. *EZHIP* non-synonymous mutations have not been reported in other pediatric brain tumor types^13^, although the *EZHIP* locus is involved in rare translocation events that result in fusion transcripts in low-grade endometrial stromal sarcoma (*MBTD1-EZHIP*^14^) and in astroblastoma (*EWSR1-EZHIP*^15^). The rare PFA cases that do not express EZHIP usually have mutations in a gene encoding the histone 3 variant H3.3, resulting in a lysine to methionine substitution at residue 27 (H3.3K27M)^16^. K27M is a mutation characteristic of a subgroup of pediatric high-grade gliomas (pHGG) defined as diffuse midline gliomas (DMGs)^17–20^. Conversely, there have been reports of rare histone 3 wild type DMG cases with expression of EZHIP^21^. The mutual exclusivity of the K27M mutation and EZHIP expression supports the idea that H3.3K27M and EZHIP function redundantly to interact and inhibit the methyl-transferase function of PRC2. Consequently, both PFA and DMGs exhibit global reduction in H3K27me3 levels that are detectable by immunohistochemistry. Therefore, although PFA and DMGs are clearly different diseases, they do share some molecular features, as was previously noted^22^.

With the present work, our goal is to describe the mutational landscape of *EZHIP* in pediatric brain tumors and to better understand the function of this gene using relevant human neural models.

## RESULTS

### Mutational analysis of EZHIP in pediatric brain tumors

The first report on *EZHIP* mutations in PFA samples in 2018 did not identify mutations in this gene in pHGG and medulloblastoma^13^. However, since then more tumor genomes have become available, providing an opportunity to revisit the mutational status of this gene across tumor types. We interrogated the Pediatric Brain Tumor Atlas (PBTA^23^, provisional cohort), which has data for 3,259 samples from a wide spectrum of central nervous system malignancies. 0.8% of patients had coding mutations in *EZHIP* (**Fig 1A**). To provide context, we also queried the mutation frequencies of other genes that are frequently mutated in pediatric brain tumor cohort, including *H3-3A* (previously known as *H3F3A*, encoding the histone 3 variant H3.3), which is mutated in 10% of individuals, *TP53* (14%) and *ATRX* (6%). Given that EZHIP is known to interact with and inhibit the catalytic function of PRC2, we also looked at genes that code for members of this complex and found that each one of them is mutated in ≤1% of patients (**Fig 1A**). Mutations in *EZHIP* tended to co-occur with mutations in *JARID2* (Benjamini-Hochberg q < 0.001), *ATRX* (q < 0.001), *EED* (q = 0.004), *TP53* (q = 0.007) and *EZH2* (q = 0.010) (**Fig 1B**). EZHIP lacks clearly defined protein domains and is predicted to be a largely disordered protein. However, a previous publication reported a mutational hotspot in a relatively ordered region at the N-terminus of the protein^13^. Using the large PBTA cohort, we found that EZHIP mutations affected amino acids throughout the protein and lacked obvious hotspots (**Fig 1C**). Regions of EZHIP where amino acid substitutions are predicted to have high pathogenic scores based on AlphaFold^24–26^ modeling tended to be free of mutations (**Fig 1D**). To further explore the mutational patterns of EZHIP, we queried databases that include adult tumors and large non-cancer cohorts. 1.3% of samples in the TCGA PanCancer Atlas^27^ (n = 10,967) had EZHIP mutations (**Fig S1A**). Amino acid substitutions occurred throughout the protein, including at regions predicted to have high pathogenic scores (**Fig S1B**). The most frequently mutated residue was E108K/D (n = 6 samples). Virtually all amino acids in EZHIP are mutated in a large cohort (n > 700,000) collected by the Genome Aggregation Database (GnomAD^28^), with R180 having the highest frequency (∼0.003 allelic frequency; **Fig S1C; Table S1**). Given the predicted disordered structure of EZHIP, and the localization of mutations in regions predicted to have low pathogenic score in cancer samples and in non-cancer cohorts, our results do not support a pathogenic role for the mutations we identified in pediatric brain tumor patients. However, future functional studies should further test this possibility.

**Figure 1.**
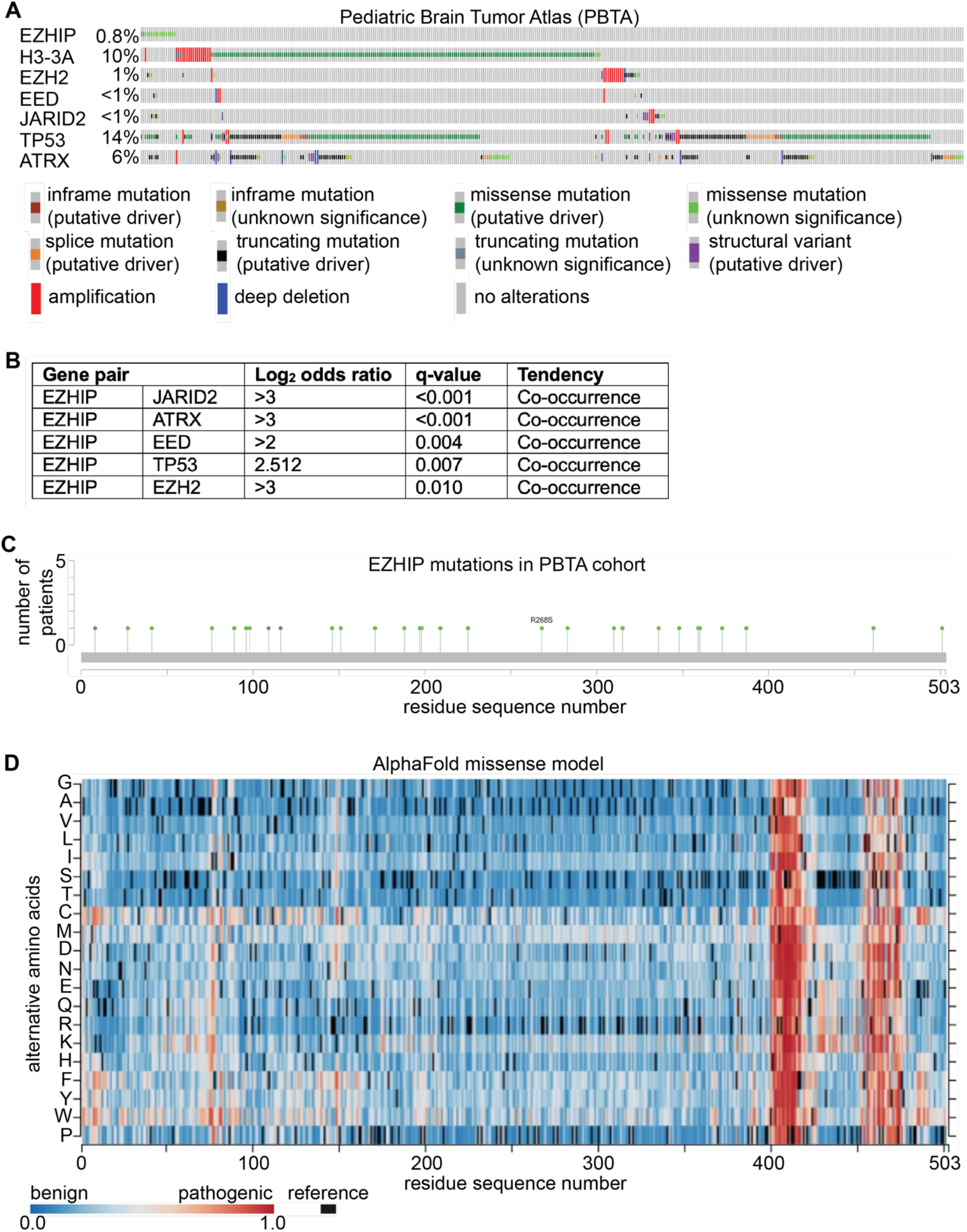
EZHIP mutations in pediatric brain tumors. **(A)** Oncoplot of EZHIP and other relevant mutations in the PBTA cohort (3,259 samples). **(B)** EZHIP mutations tend to co-occur with mutations in TP53 and genes encoding PRC2 members. **(C)** EZHIP mutations in pediatric brain tumors are patient-specific. **(D)** EZHIP mutations affect residues not predicted to be deleterious or pathogenic.

### EZHIP is mutated in non-PFA pediatric brain tumors

Next, we looked at the diagnoses of the pediatric brain tumor patients with EZHIP mutations. Five patients were diagnosed with ependymoma, 3 of which corresponded to the PFA subtype, as expected (**Fig 2A**). The average age of the ependymoma patients with EZHIP mutations was 6 years and the median age was 3 years. Interestingly, 3 other patients were diagnosed with medulloblastoma, including 2 SHH and one WNT cases, with an average age of 4 years and a median age of 3 years. Finally, 4 patients were diagnosed with HGG, with an average age of 5.2 years and a median age of 5 years. Two of the HGG cases were molecularly defined as DMG, H3K27 altered, and the remaining 3 were histone 3 wild-type. To determine the robustness of the mutational calls in *EZHIP*, we evaluated the quality of each read that supported the calls and found them of high quality (**Fig 2B**). Additionally, the *EZHIP* mutations were clearly somatic and were not found in the blood samples of the patients (**Fig 2B**). The identification of EZHIP mutations in HGG patients is intriguing because (i) they were not known to occur and (ii) it was previously suggested that DMG and PFA exist along a continuum of molecular features that may result in shared genetic, metabolic and epigenetic dependencies^22^. We therefore decided to perform more in-depth analyses of the mutational landscapes of EZHIP-mutated HGG. Coding mutations in only one gene – *EGFR* – co-occurred with *EZHIP* mutations in all HGG samples (**Fig 2C**). 3 of 5 samples had mutations in *TP53*, while two HGG samples had mutations in *RYR1/2/3*, encoding a family of ryanodine receptors with roles in intracellular Ca^2+^ signaling, and *KMT2D*, which codes for a histone methyltransferase with critical roles in maintaining enhancer elements in mammalian cells. Two *EGFR* mutations occur at known cancer hotspots and are likely to be drivers (**Fig 2D**). A full list of mutations that co-occur with *EZHIP* mutations in HGG cases in the PBTA cohort can be found in **Table S2**. The key findings that *EZHIP* is mutated in pediatric HGG samples and that *EZHIP* mutations co-occur with *EGFR* mutations was validated in an independent cohort from the St Jude Research Hospital^29^ (**Fig S2A-B**). We next asked whether the mutated *EZHIP* gene was transcribed in HGG samples. RNA-seq data were available for 3 of 5 *EZHIP* mutant pHGG samples in the PBTA cohort, and *EZHIP* was transcribed in all 3 samples (**Fig S3A-C**). Interestingly, mutant *EZHIP* was also expressed in a pHGG sample (7316-9049) with an H3K27 mutation (**Fig S3C**), although it was until now assumed that *EZHIP* expression and histone 3 mutations are mutually exclusive. Taken together, our data show that *EZHIP* mutations occur not only in PFA, but also in pediatric cases of medulloblastoma and HGG, where they tend to co-occur with mutations in *EGFR*. In all tumor types, *EZHIP* mutations occur in young children.

**Figure 2.**
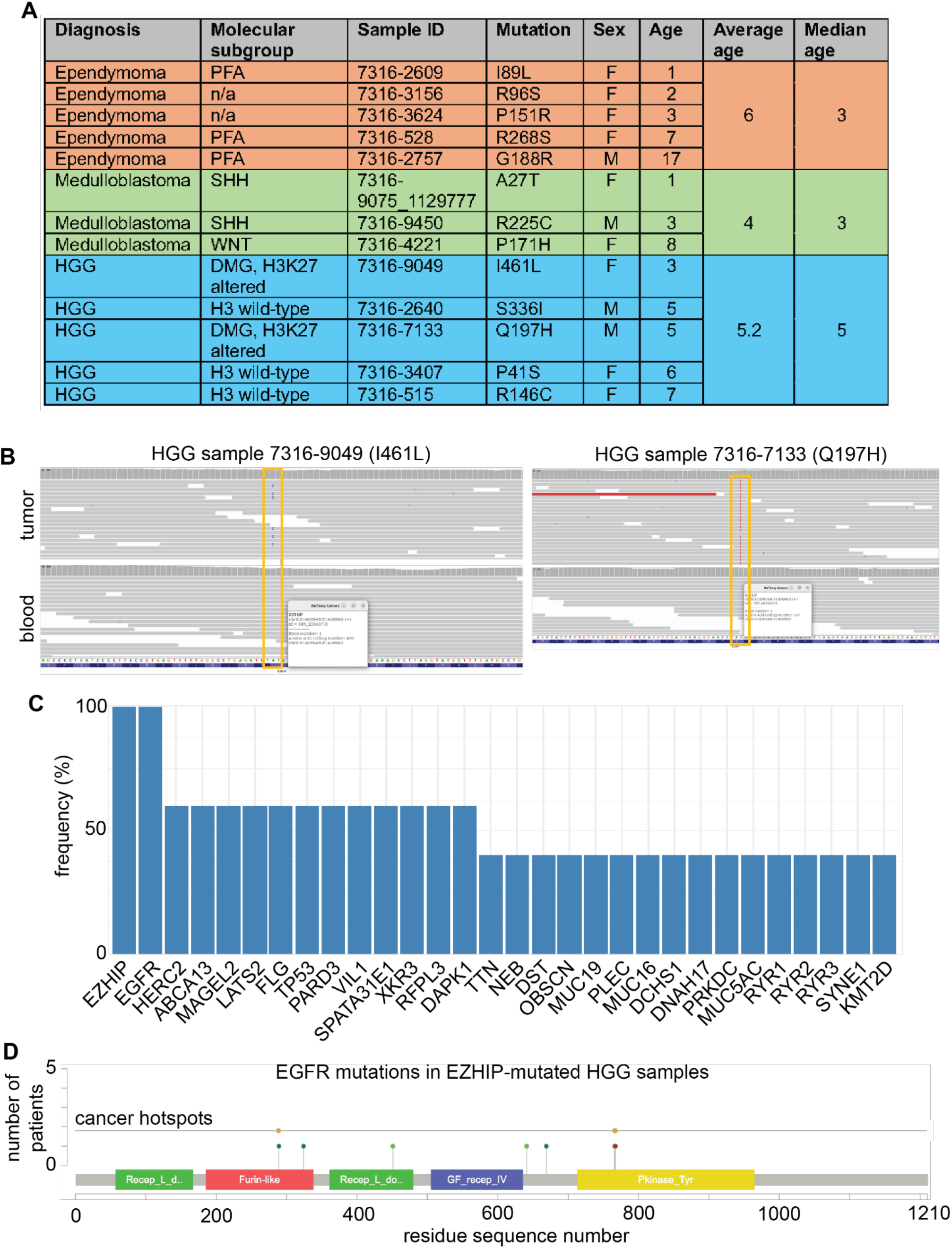
EZHIP mutations are not exclusive to PFA. **(A)** Medulloblastoma and medulloblastoma cases have EZHIP mutations. **(B)** Examples of somatic EZHIP mutations in HGG. **(C)** EZHIP mutations co-occur with EGFR lesions in HGG patients. **(D)** Some of the EGFR mutations in EZHIP-mutant HGG cases are known to be pathogenic.

### EZHIP expression activates a neuronal expression program

Surprisingly for a protein that may have important roles in pediatric brain tumors, very little information is currently available on the function of EZHIP in relevant neural models. Our analyses of previously published data from prenatal human samples (first and second trimester post-conception) indicate that *EZHIP* is not expressed in any pre-natal brain cell types for which single-cell RNA-seq data were available^30,31^ (**Fig S4A-B**). However, it may be possible that *EZHIP* is expressed at low levels in rare pre-natal cell types that were not captured in the datasets we analyzed. As a step toward understanding the role of EZHIP in neural systems, we used a PiggyBac transposition system to constitutively express this protein (EZHIP OE) in human neural progenitor cell (hNPC) models and that we described in a previous publication^10^. We used these human neural models to investigate the transcriptional changes associated with EZHIP expression. Two hNPC cultures expressing EZHIP (two biological replicates for each), and their respective controls, were used for transcriptomic analysis by bulk RNA-seq (see Methods). Differential gene expression analysis was performed using RSEM expected counts as input for the edgeR/limma^32^ pipeline. The analysis identified 3,503 and 4,120 upregulated genes, and 3,691 and 4,143 downregulated genes in hNPC5 and hNPC6, respectively. Of these, 1,426 genes were upregulated in both models, and 1,384 genes were downregulated in both EZHIP OE hNPCs (**Fig 3A, Fig S4C**). Gene Set Enrichment Analysis (GSEA)^33^ showed that several gene sets associated with synaptic function were enriched in EZHIP OE hNPCs compared to controls. These gene sets were “protein-protein interactions at synapses” (**Fig 3B**), “neurotransmitter release cycle” (**Fig 3C**), “neurexins and neuroligins” (**Fig 3D**), and “epithelial mesenchymal transition” (**Fig S4D**). Similarly, Gene Ontology (GO) analysis of RNA sequencing data from EZHIP-expressing hNPCs revealed significant enrichment in processes related to neuron cell-cell adhesion, synaptic membrane adhesion, and positive regulation of excitatory postsynaptic potential (**Fig S4E**). These results highlight a potential role for EZHIP in neural-like signaling.

**Figure 3.**
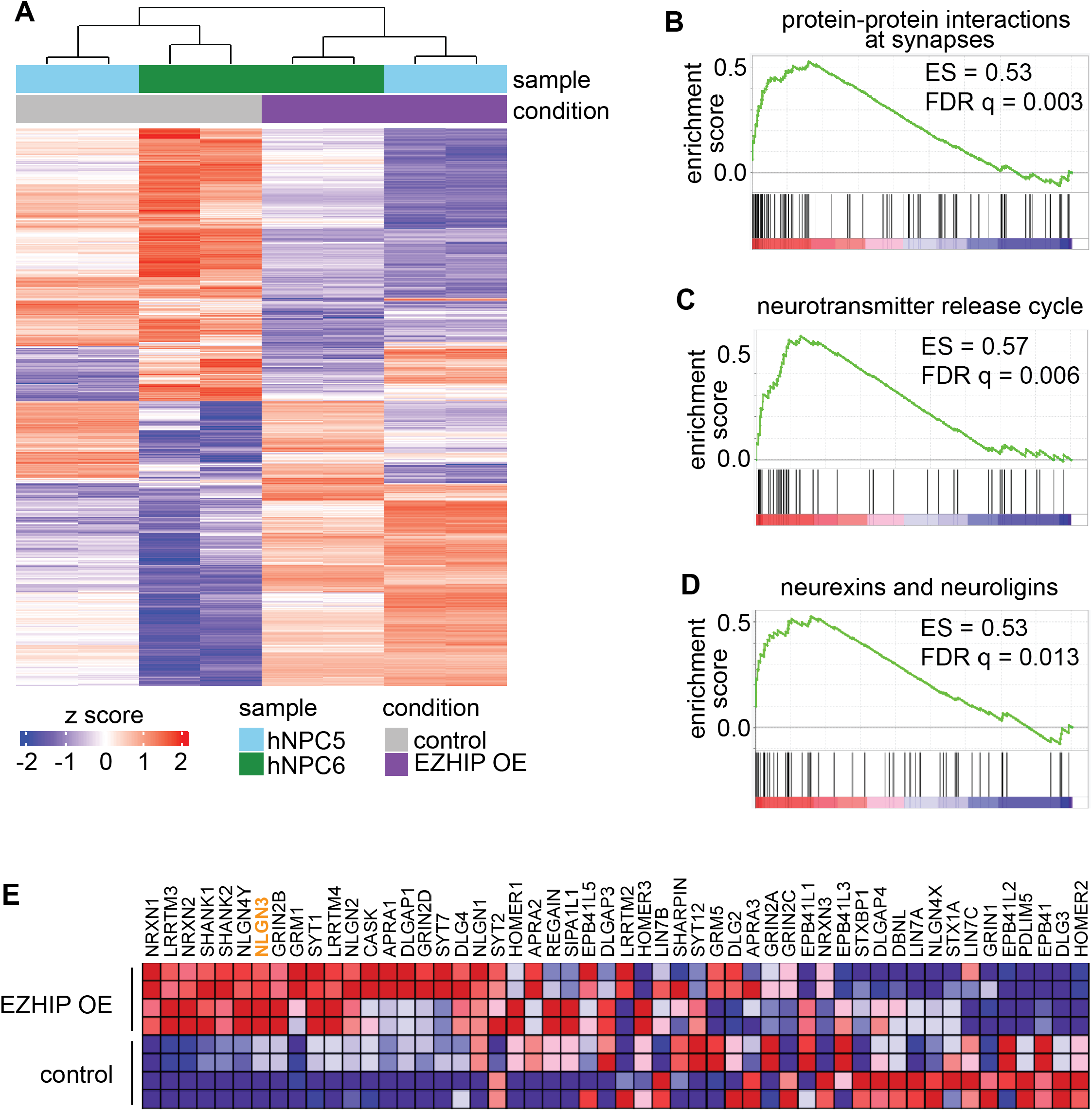
Effects of expression of EZHIP in human neural models. **(A)** RNA-seq of two independent hNPC models that over-express EZHIP. **(B-D)** EZHIP expression is associated with enrichment of gene signatures related to synaptic signaling. **(E)** EZHIP expression potentiates expression of neurexins and neuroligins.

The association of EZHIP expression with transcriptional programs of neuro-synapse function was interesting because synapses among tumor cells and between tumor cells and neurons have been shown to positively affect malignant growth in pediatric and adult HGG^34–37^. In HGG, the neuroligin NLGN3 has distinctive pro-tumor effects^35^. Among the genes in the “neurexin and neuroligin” gene set, *NLGN3* is strongly and consistently expressed in all biological replicates in EZHIP OE hNPCs (**Fig 3E**). To validate the transcriptomic data, we performed western blots for NLGN3 in 3 independent hNPC models and found that this protein was expressed at higher levels in EZHIP OE hNPCs than in their respective controls (**Fig 4A**).

**Figure 4.**
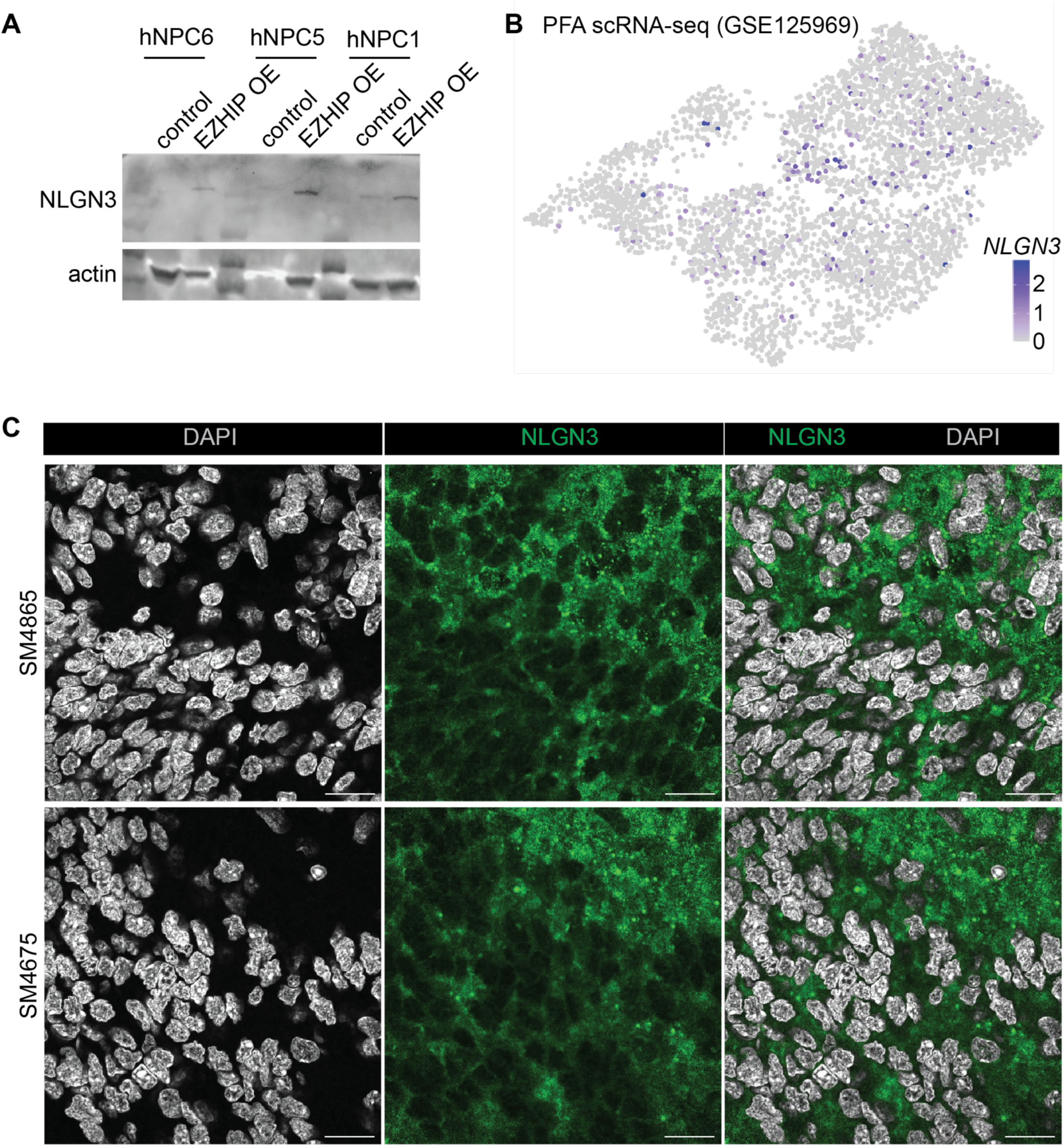
NLGN3 is produced by PFA tumors. **(A)** Western blot validates production of NLGN3 in EZHIP-expressing hNPCs. **(B)** *NLGN3* is expressed in PFA cells. scRNA-seq from Gillen et al. **(C)** Immunofluorescence for NLGN3 on sections from two independent PFA tumors. Scale bar: 20 μm.

To test if our findings could be generalized to PFA, which express EZHIP, we re-analyzed a previously published scRNA-seq dataset^38^ generated from surgical specimens. The results show that some PFA cells express *NLGN3* (**Fig 4B**), supporting the conclusion of our bulk RNA-seq results. Next, we investigated whether NLGN3 is detectable at the protein level in PFA. Immunofluorescence experiments showed the presence of NLGN3 in two independent PFA surgical specimens (SM4865 and SM4675; **Figs 4C, S4F**).

Overall, our analyses and experiments in human neural models underscore the previously unknown association between EZHIP expression and modulation of genes involved in synaptic function and neural connectivity.

### EZHIP expression affects major metabolic pathways

While no recurrent PFA alterations aside from EZHIP have been identified, PFA tumors exhibit distinct metabolic gene signatures that reduce histone methylation and enhance histone demethylation. This suggests that EZHIP may be linked to metabolic regulation through its influence on key gene signatures. To identify disease-relevant metabolic pathways in PFA, an untargeted metabolomics analysis was conducted using PFA patient tumors and normal human brain tissue (Michaelraj et al., 2020). Upregulated pathways in PFA include purine metabolism, glycerophospholipid metabolism, and arginine biosynthesis (**Figure 5A**). In contrast, downregulated pathways included glycine, serine, and threonine metabolism, one carbon pool by folate, as well as phenylalanine, tyrosine, and tryptophan metabolism (**Figure 5B**). After establishing a PFA-specific steady-state metabolic profile, the impact of EZHIP expression on metabolism was investigated. Metabolomic analysis was performed on EZHIP-expressing hNPCs compared to controls using liquid chromatography high resolution mass spectrometry (LC-HRMS) (**Figure 5C, Table S3**). EZHIP expression indeed led to profound alterations in amino acid metabolism, glycolysis, and the Krebs/TCA cycle, revealing a previously unrecognized link between this protein and metabolic reprogramming (**Figure 5D-F**). Integration of transcriptomic and metabolic profiles uncovered an unrecognized role of EZHIP in rewiring key regulatory pathways involved in cellular metabolism and stress responses, including HIF-1α signaling, phosphatidylinositol signaling, and one-carbon pool by folate (**Figure 6A**). When studying metabolic pathways specifically, glycerophospholipid metabolism, glycolysis/gluconeogenesis, and one carbon pool by folate were impacted by EZHIP expression (**Figure 6B**). One-carbon metabolism, which includes methionine and folate metabolism, is essential for the synthesis of polyamines and other cellular building blocks. Given that folate-driven one-carbon metabolism was identified across all transcriptomic and metabolic pathways, RNA sequencing analysis was employed, revealing a downregulation of polyamine metabolism. (**Figure 6C**). Key metabolites and enzymes involved in both methionine and polyamine metabolism were collectively decreased (**Figure 6D**). Of interest, AMD1 – an enzyme responsible for generating decarboxylated S-adenosylmethionine (dcSAM) within the methionine metabolism pathway - was notably reduced. Given the role of dcSAM in supporting the production of polyamines such as spermidine and spermine, our findings suggest that EZHIP expression in neural models may suppress polyamine biosynthesis through coordinated transcriptional and metabolic repression of one-carbon and methionine cycle intermediates.

**Figure 5.**
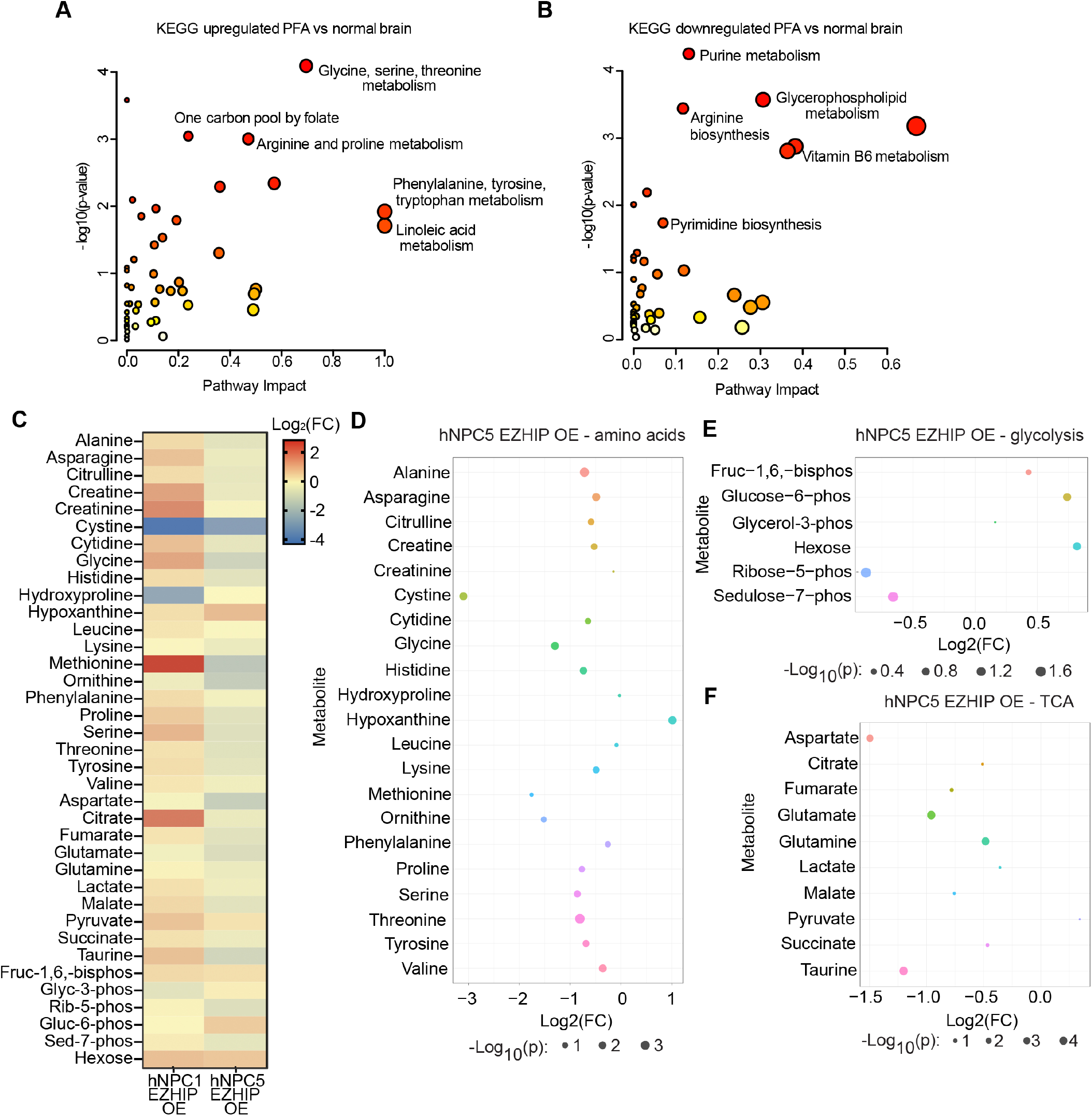
Targeted metabolomics analysis. **(A)** KEGG upregulated pathway analysis of PFA (n=15) and normal brain (n=5) from Michaelraj et al. Cutoff included metabolites >1 fold change. **(B)** KEGG downregulated pathway analysis of PFA (n=15) and normal brain (n=5) from Michaelraj et al. Cutoff included metabolites <-1 fold change. **(C)** Heatmap showing log_2_FC in metabolite alterations in EZHIP overexpression versus control in hNPC11 and hNPC5 cell lines. **(D)** Bubble plot showing amino acid alterations in hNPC5 EZHIP overexpression compared to control. **(E)** Bubble plot showing glycolysis metabolite alterations in hNPC5 EZHIP overexpression compared to control. **(F)** Bubble plot showing TCA/Kreb’s cycle metabolite alterations in hNPC5 EZHIP overexpression compared to control.

**Figure 6.**
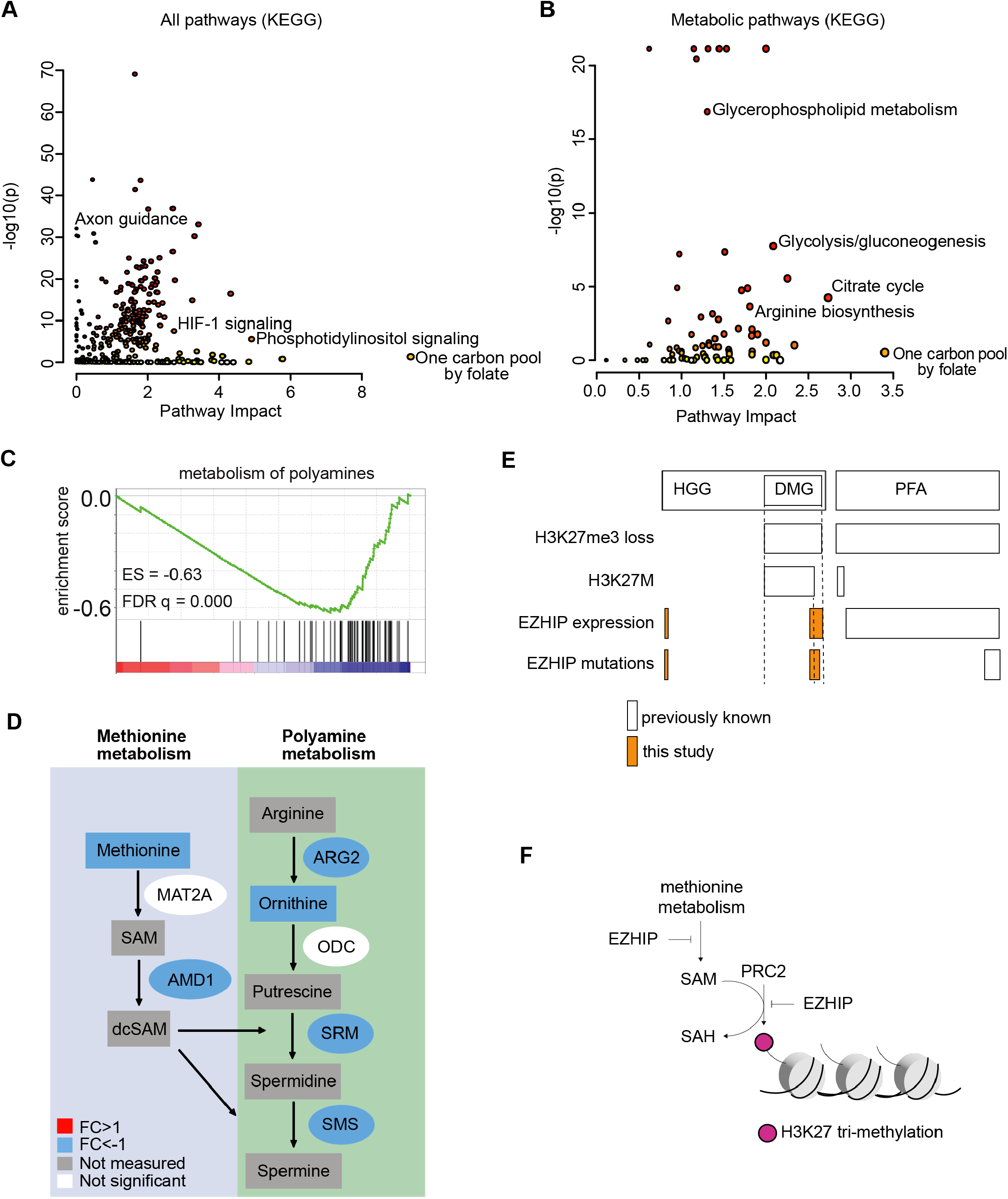
Combined analysis of RNA sequencing and targeted metabolomics. **(A)** KEGG all pathway integrated analysis of HFT5 RNA sequencing and targeted metabolomics using fold change. Bubble size indicates pathway impact. **(B)** KEGG metabolism pathways analysis of HFT5 RNA sequencing and targeted metabolomics using fold change. Bubble size indicates pathway impact. **(C)** RNA-sequencing supports negative enrichment of genes associated with polyamine metabolism. **(D)** Specific analysis of polyamine metabolism, where downregulated genes and metabolites are shown in blue. **(E)** Diagram summarizing the mutational patterns of EZHIP in high-grade glioma (HGG) and PFA ependymoma, highlighting the new findings of this study. **(F)** Cartoon to illustrate a proposed model for the global loss of H3K27me3 observed in PFA ependymoma.

## DISCUSSION

There currently is a widespread assumption that *EZHIP* is mutated exclusively in PFA, and that its expression is mutually exclusive with the presence of the K27M histone 3 mutation in both PFA and DMG. However, here we demonstrate that *EZHIP* mutations occur in other types of brain tumors at low frequency. For instance, we found HGG cases (including histone 3-altered DMG) and medulloblastoma cases with somatic *EZHIP* mutations. Of note, *EZHIP* and *EGFR* mutations co-occurred in all cases of pediatric HGG that we identified in two separate cohorts. The co-occurrence of *EZHIP* and *EGFR* mutations is intriguing and may point to some aspects of the biology of HGG and EZHIP function that are not currently understood and deserve further investigation. Although the number of cases was small, we also noticed that pediatric HGG patients with *EZHIP* mutations tended to be young (median age of 5 years), further suggesting underlying biological processes specific to this population. In addition, it is also presumed that *EZHIP* is not expressed in H3-altered DMG. However, we now provide evidence that *EZHIP* is in fact expressed in this molecularly-defined tumor type (**Figure 6E**).

Previous functional studies on EZHIP have focused on its direct inhibition of PRC2. Here, we used human neural models to further investigate the relationship between EZHIP and regulation of gene expression. We found that EZHIP expression has a strong effect on activating genes involved in neuronal synapse function. One of the key classes of upregulated genes are neurexins and neuroligins, including *NLGN3*, which has been investigated as a mediator of cell communication between neurons and glioma cells^36^. Immunohistochemistry experiments using surgical specimens confirmed expression of NLGN3 in PFA cells. This result is interesting because previous studies on HGG determined that expression of NLGN3 occurred in non-neoplastic neurons adjacent to the malignant cells^35^. It is therefore possible that PFA cells can express NLGN3 to boost intrinsic neuronal-like programs of cell communication that are beneficial for tumor growth. Future studies will be needed to explore the role of neuronal-like communication in PFA.

A large fraction of genes was downregulated in EZHIP-expressing hNPCs. A significant fraction of these genes is involved in regulation of metabolism. We therefore performed metabolomics on EZHIP-expressing and control hNPCs, and integration of metabolomics and transcriptomics data identified significant reduction in metabolism of methionine and polyamines. The levels of methionine itself were significantly decreased in EZHIP-expressing models. Methionine is required for the production of SAM, the universal methyl group donor required for histone methylation. It is therefore possible that the global loss of H3K27me3 that is typical of PFA might be a consequence of EZHIP-mediated repression of methionine metabolism, in addition to the direct inhibition of PRC2 by EZHIP (**Figure 6F**). Expression of EZHIP might therefore contribute to both DNA and histone hypomethylation observed in PFA.

Methionine and polyamine metabolism share intermediates and we found that both pathways were repressed in EZHIP-expressing hNPCs. Further, a gene that mediates polyamine metabolism, *AMD1*, is significantly downregulated in PFA compared to other pediatric brain tumors, indicating that decreased polyamine metabolism is a potential vulnerability in this tumor type. DMG is also characterized by decreased polyamine metabolism, which represents a potential vulnerability. Preclinical and clinical trials for DMG and pediatric HGG have tested the efficacy of inhibitors of polyamine metabolism. Multiple clinical trials are testing AMTX-1501^39^, an inhibitor of polyamine transport, in patients with CNS malignancies (NCT05717153, early phase 1; NCT06465199, phase 1/2), including DMG and pediatric HGG. Further testing of polyamine transport inhibition in preclinical models of PFA and potentially extending current clinical trials to PFA patients might be a potential translation opportunity.

## MATERIALS AND METHODS

### Mutation analyses in pediatric and adult cohorts

Mutational status for selected genes in pediatric brain tumor patients was determined by accessing provisional data from the Pediatric Brain Tumor Atlas (PBTA) through pedcBioPortal (http://pedcbioportal.kidsfirstdrc.org). Oncoplots, mutation co-occurrence statistics and lollipop plots were generated by the pedcBioPortal interactive hub. Biological sex of samples, age of the patient at diagnosis and other sample information was identified using the clinical information tab on the portal. To confirm *EZHIP* mutation status, we downloaded sequencing files for each sample available through CAVATICA (https://pgc-accounts.sbgenomics.com) through the Children’s Brain Tumor Network (CBTN). BAM files were opened in IGV and inspected for sequencing read quality at each mutation. GnomAD (v4.1.0) data were accessed at https://gnomad.broadinstitute.org.

### AlphaFold pathogenic scores

Pathogenic scores for amino acid substitution in the EZHIP protein were accessed through the AlphaFold website (https://alphafold.ebi.ac.uk).

### RNA extraction and bulk RNA-seq

RNA was extracted from *EZHIP* OE hNPCs and their respective controls using the RNEasy Mini kit (79254, Qiagen) as per manufacturer instructions. Libraries were constructed at the Center for Health Genomics and Informatics at the University of Calgary Cumming School of Medicine using the NEBNext Ultra II Directional RNA Library prep kit (New England Biolabs) with ribosomal RNA depletion. RNA-seq data analysis was conducted on the HPC cluster maintained by the Biostatistics and Informatics Shared Resource (BISR) at the Dan L Duncan Comprehensive Cancer Center and Baylor College of Medicine. FASTQ files were processed using the nf-core framework^40^ with the RNA-seq pipeline (version 3.14.0, https://doi.org/10.5281/zenodo.10471647).

The reference files used were Homo_sapiens.GRCh38.dna_sm.primary_assembly.fa.gz and Homo_sapiens.GRCh38.111.gtf.gz, and STAR_RSEM was employed as the aligner.

Gene counts were analyzed for differential expression using the R packages edgeR (version 4.0.16)^41^. Gene annotation was performed with the R package org.Hs.eg.db version 3.18.0, excluding duplicated Ensemble gene ID entries. The standard limma version 3.58.1 was used to complete the differential expression analysis^42^.

Gene Ontology (GO) enrichment analysis was carried out using the R package topGO (version 2.54.0, https://doi.org/10.18129/B9.bioc.topGO), applying the “elim” algorithm and “fisher” statistics with a cutoff of 0.01. Significant GO terms were defined by odds ratio > 1, weightFisher < 0.05, and number of annotations >= 10.

For Gene Set Enrichment Analysis^33,43^, expected counts were normalized using the R package DESeq2 version 1.42.1^44^. Of the original 21,507 features, 18,052 genes remained after collapsing into gene symbols. Gene set size filters were set at a minimum of 15 and a maximum of 500, resulting in 1,707 gene sets that included key BIOCARTA, HALLMARK, KEGG, REACTOME, and Pathway Interaction Database (PID) pathways. The analysis was performed with 1,000 permutations of type “gene_set.”

### Immunofluorescence of primary patient samples

Samples were deparaffinized with xylene and dehydrated, followed by antigen retrieval with EDTA buffer (1 mM EDTA pH 8.0; 0.05% Tween-20) in a pressure cooker for 35 minutes. Samples were blocked with BSA (5% BSA in PBS with 0.1% Triton-X 100) for 1 hour, then incubated with primary antibody (mouse Neuroligin 3/NLGN3 Antibody (S110-29): 1:100, Novus NBP2-42200) overnight at 4°C in staining buffer. Secondary antibody staining was performed for 1 hour at room temperature using fluorescently conjugated secondary antibodies:(Goat anti-Mouse IgG (H+L) Cross-Adsorbed Secondary Antibody, Alexa Fluor™ 488 (dilution 1:500; A-11001, Thermo Fisher). Incubated for 5 minutes in DAPI (dilution 1:1000; 62248, Thermo Fisher). Washes were performed using PBS with 0.1% Tween and mounted with Prolong Diamond antifade (P36965, Life Technologies). Images were acquired using the ZEISS LSM 880 Airyscan confocal microscope.

### Western blots

Protein concentration of samples was determined using the DC (detergent compatible) protein assay (5000112, Bio-Rad). Samples were prepared in a total volume of 20 μL at 10 μg/μL in Laemmli loading buffer. Samples were run on 10% Mini-PROTEAN gels (4568044, Bio-Rad). Primary antibodies used: Rabbit Neuroligin 3/NLGN3 Antibody (0.04 μg/mL, Novus NBP1-90080). Mouse monoclonal anti-β-ACTIN (Sigma-Aldrich, A5441, Lot# 127M4866V, Clone AC-15) was used at 1:1000 dilution. Secondary antibodies used: Goat anti-rabbit IgG H&L (hrp) (ab6721, Abcam) and Goat anti-mouse IgG H&L (hrp) (ab6789) at 1:20,000 dilutions.

### Untargeted high-resolution LC-HRMS

#### Sample preparation

Metabolic quenching and polar metabolite pool extraction was performed by adding ice cold 80% methanol (aqueous) at a ratio of 500 µL buffer per 1e6 cells. Deuterated (D_3_)-creatinine and (D_3_)-alanine, (D_4_)-taurine and (D_3_)-lactate (Sigma-Aldrich) was added to the sample lysates as an internal standard for a final concentration of 10 µM. Samples are scraped into Eppendorf tubes on ice, homogenized using a 25°C water bath sonicator and the supernatant was then cleared of protein by centrifugation at 16,000xg. 2 µL of cleared supernatant was subjected to online LC-MS analysis.

#### LC-HRMS Method

Analyses were performed by untargeted LC-HRMS. Briefly, Samples were injected via a Thermo Vanquish UHPLC and separated over a reversed phase Thermo HyperCarb porous graphite column (2.1×100 mm, 3 μm particle size) maintained at 55°C. For the 20 minute LC gradient, the mobile phase consisted of the following: solvent A (water / 0.1% FA) and solvent B (ACN / 0.1% FA). The gradient was the following: 0-1 minute 1% B, increase to 15% B over 5 minutes, continue increasing to 98%B over 5 minutes, hold at 98%B for five minutes, reequillibrate at 1%B for five minutes. The Thermo IDX tribrid mass spectrometer was operated in both positive and negative ion mode, scanning in ddMS^2^ mode (2 μscans) from 70 to 800 m/z at 120,000 resolution with an AGC target of 2e5 for full scan, 2e4 for ms^2^ scans using HCD fragmentation at stepped 15,35,50 collision energies. Source ionization setting was 3.5 and 2.4kV spray voltage respectively for positive and negative mode. Source gas parameters were 35 sheath gas, 12 auxiliary gas at 320°C, and 8 sweep gas. Calibration was performed prior to analysis using the Pierce™ FlexMix Ion Calibration Solutions (Thermo Fisher Scientific). Integrated peak areas were then extracted manually using Quan Browser (Thermo Fisher Xcalibur ver. 2.7). Purified standards were then purchased and compared in retention time, m/z, along with ms^2^ fragmentation patterns to validate the identity of significant hits.

### Metabolic pathway analysis and integrated transcriptomics and metabolic profiles

Pathway enrichment of untargeted metabolomics of patient samples^12^, was completed using Metaboanalyst 6.0 via pathway enrichment, and metabolites exhibiting a ≥1-fold change with p < 0.05 were considered significant. Integration of bulk transcriptomics and targeted metabolic profiles was completed using Metaboanalyst 6.0 via the Joint Pathway Analysis module. Fold change for each gene and metabolite was uploaded, with a cutoff of p<0.05 and FC=2. Pathway analysis for integrated metabolic pathways and all pathways integrated were visualized using hypergeometric test for enrichment analysis, degree centrality for topology measure, and combined queries for integration methods.

## DATA AVAILABILITY

RNA-seq data were deposited to the Gene Expression Omnibus (GEO) under accession number GSE291006.

## ACKNOWLEDGEMENTS

This work was supported by Canadian Institutes of Health Research (CIHR) project grants (PJT-156278 and PJT-173475) and a Canada Research Chair to MG.

## AUTHOR CONTRIBUTIONS

EH, HMC, TAG, STY, GAN, FJ and HM performed the experiments described in this manuscript. MG, SA, MDT and WY supervised and planned experimental activities. EH, HMC, TAG, SA and MG contributed to writing the manuscript. All authors contributed comments and edits.

## Notes

### Competing Interest Statement

The authors have declared no competing interest.

