## Supplemental figures for "EZHIP boosts neuronal-like synaptic gene programs and depresses polyamine metabolism"

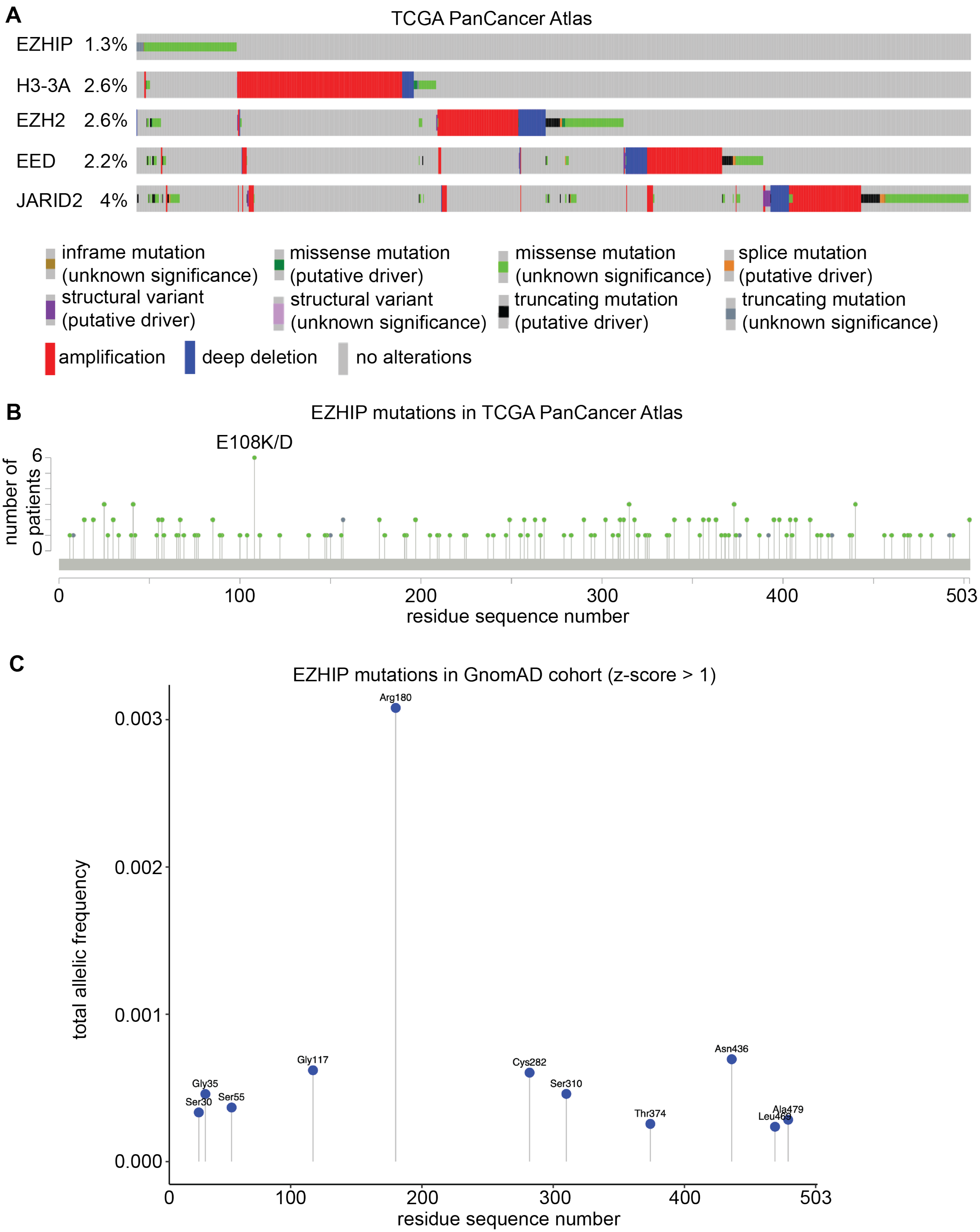
A
TCGA PanCancer Atlas
EZHIP 1.3%
H3-3A 2.6%
EZH2 2.6%
EED 2.2%
JARID2 4%
inframe mutation (unknown significance)
missense mutation (putative driver)
missense mutation (unknown significance)
splice mutation (putative driver)
structural variant (putative driver)
structural variant (unknown significance)
truncating mutation (putative driver)
truncating mutation (unknown significance)
amplification
deep deletion
no alterations
B
EZHIP mutations in TCGA PanCancer Atlas
E108K/D
number of patients
6
0
0
100
200
300
400
503
residue sequence number
C
EZHIP mutations in GnomAD cohort (z-score > 1)
total allelic frequency
0.003
0.002
0.001
0.000
Arg180
Gly117
Gly35
Ser30
Ser55
Cys282
Ser310
Thr374
Asn436
Ala479
0
100
200
300
400
503
residue sequence number

**Figure S1. EZHIP mutations across pediatric brain tumors, adult cancers, and general population cohorts.**
**(A)** Oncoplot illustrating mutations in *EZHIP* and other genes in cancer samples of different types and etiology collected by the TCGA.
**(B)** Lollipop plot indicating EZHIP residues that are mutated in tumor samples collected by the TCGA.
**(C)** Lollipop plot indicating the distribution and frequency of mutated EZHIP residues in the general population (GnomAD cohort).

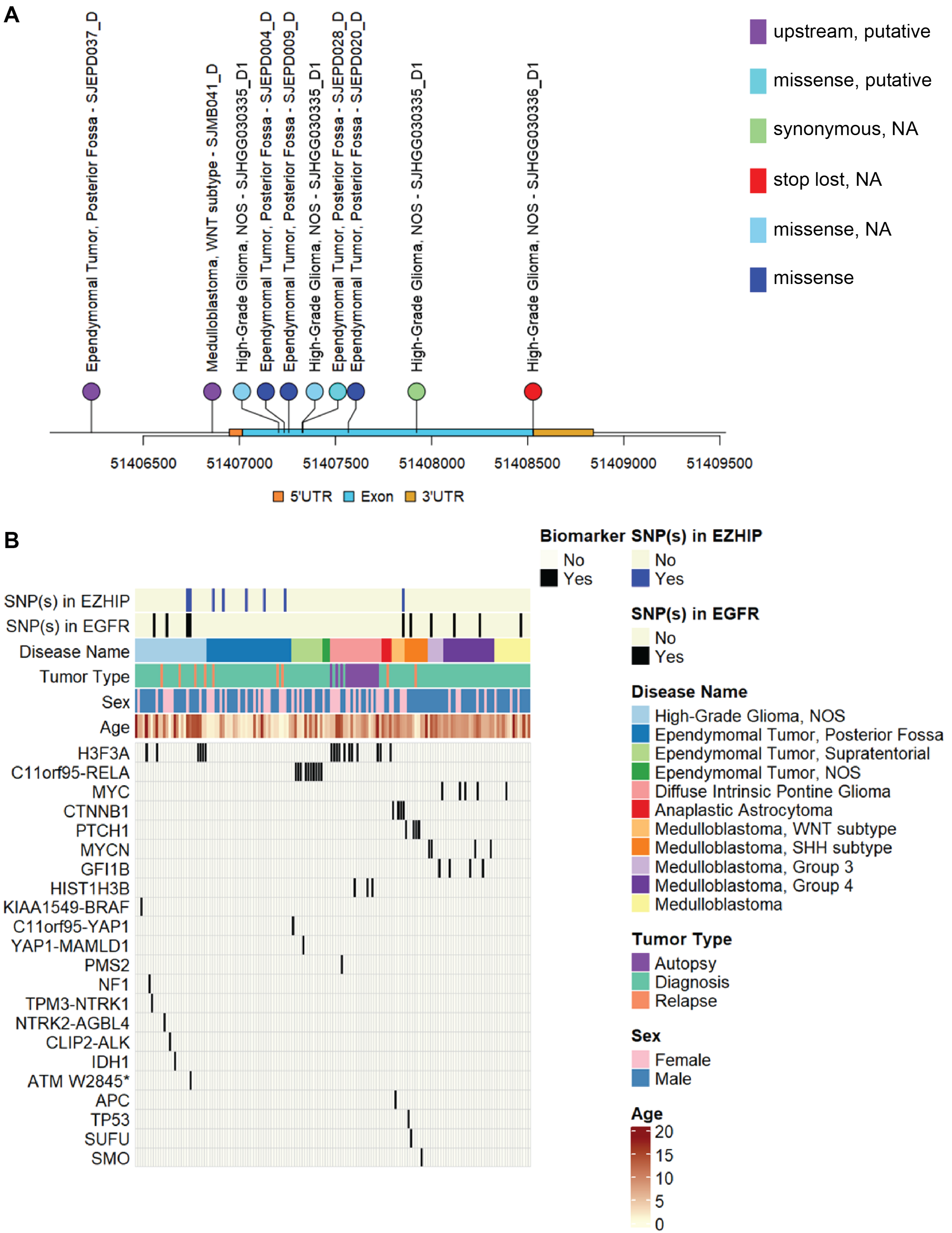
A
Ependymomal Tumor, Posterior Fossa - SJEPD037_D
Medulloblastoma, WNT subtype - SJMB041_D
High-Grade Glioma, NOS - SJHGG030335_D1
Ependymomal Tumor, Posterior Fossa - SJEPD004_D
Ependymomal Tumor, Posterior Fossa - SJEPD009_D
High-Grade Glioma, NOS - SJHGG030335_D1
Ependymomal Tumor, Posterior Fossa - SJEPD028_D
Ependymomal Tumor, Posterior Fossa - SJEPD020_D
High-Grade Glioma, NOS - SJHGG030335_D1
High-Grade Glioma, NOS - SJHGG030336_D1
upstream, putative
missense, putative
synonymous, NA
stop lost, NA
missense, NA
missense
51406500
51407000
51407500
51408000
51408500
51409000
51409500
5'UTR
Exon
3'UTR
B
SNP(s) in EZHIP
SNP(s) in EGFR
Disease Name
Tumor Type
Sex
Age
H3F3A
C11orf95-RELA
MYC
CTNNB1
PTCH1
MYCN
GFI1B
HIST1H3B
KIAA1549-BRAF
C11orf95-YAP1
YAP1-MAMLD1
PMS2
NF1
TPM3-NTRK1
NTRK2-AGBL4
CLIP2-ALK
IDH1
ATM W2845*
APC
TP53
SUFU
SMO
Biomarker
No
Yes
SNP(s) in EZHIP
No
Yes
SNP(s) in EGFR
No
Yes
Disease Name
High-Grade Glioma, NOS
Ependymomal Tumor, Posterior Fossa
Ependymomal Tumor, Supratentorial
Ependymomal Tumor, NOS
Diffuse Intrinsic Pontine Glioma
Anaplastic Astrocytoma
Medulloblastoma, WNT subtype
Medulloblastoma, SHH subtype
Medulloblastoma, Group 3
Medulloblastoma, Group 4
Medulloblastoma
Tumor Type
Autopsy
Diagnosis
Relapse
Sex
Female
Male
Age
20
15
10
5
0

**Figure S2.** ***EZHIP*** **mutations in pediatric brain tumor samples in the St Jude cohort.**
**(A)** Lollipop plot illustrating the distribution of mutations at the *EZHIP* locus in brain tumor samples included in the St Jude cohort.
**(B)** Oncoplot representing the co-occurrence of mutations in *EZHIP* and other genes in brain tumor samples that are included in the St Jude cohort.

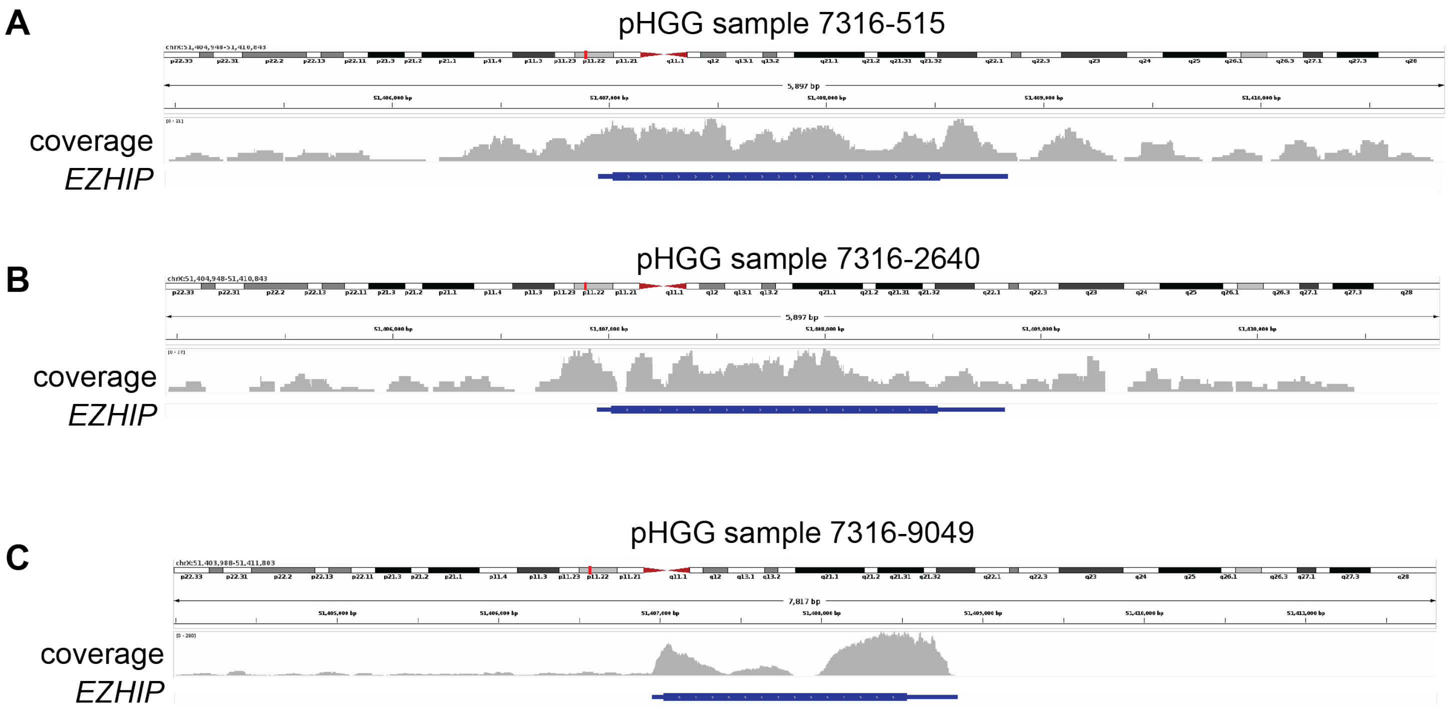


**Figure S3. Transcription of *EZHIP* in pHGG samples.**
**(A)** IGV plot displaying RNA-seq read coverage for pHGG sample 7316-515 from the CBTN cohort. Read range: [0, 21].
**(B)** IGV plot displaying RNA-seq read coverage for pHGG sample 7316-2640 from the CBTN cohort. Read range: [0, 17].
**(C)** IGV plot displaying RNA-seq read coverage for pHGG sample 7316-9049 from the CBTN cohort. Read range: [0, 280].

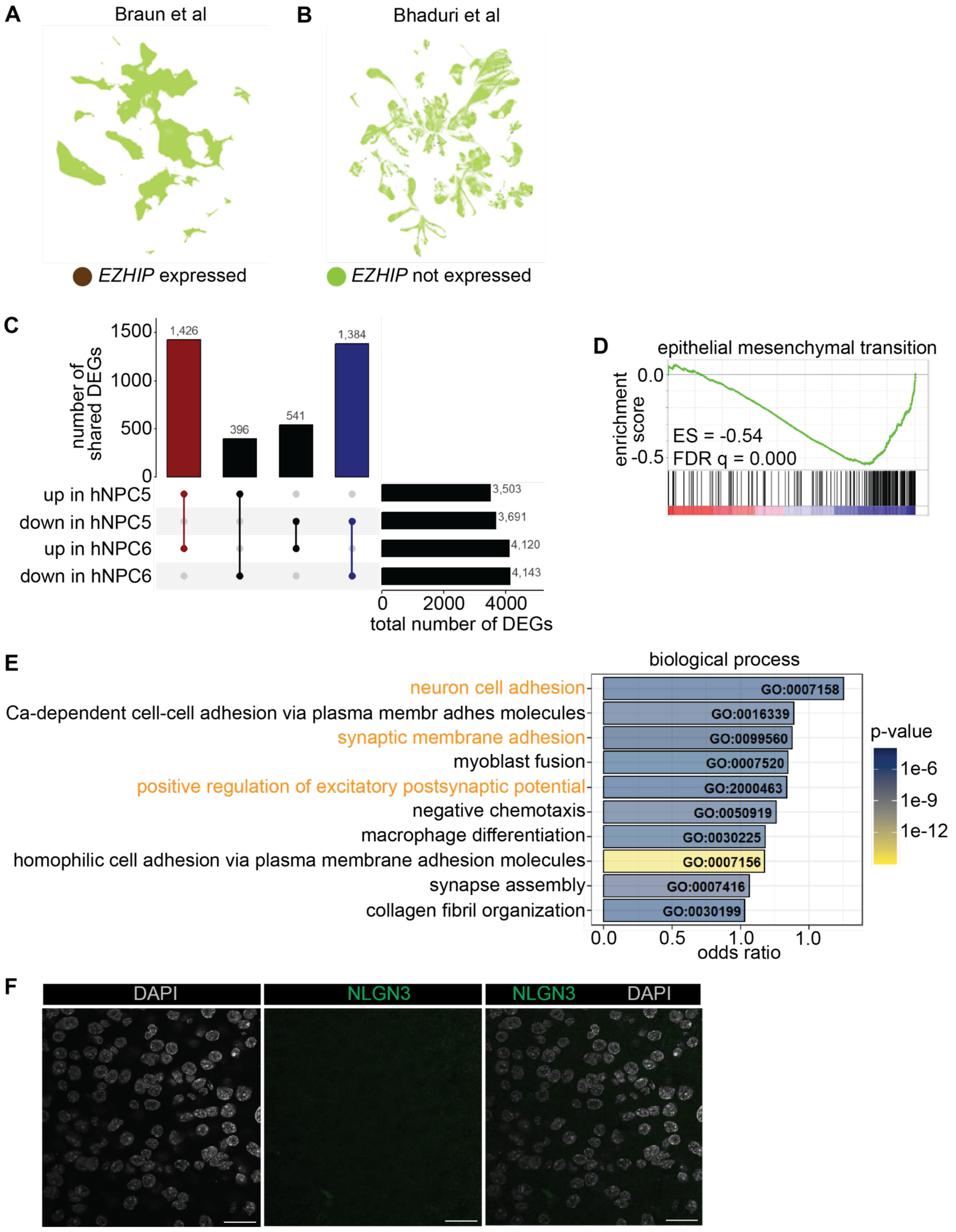
A
Braun et al
B
Bhaduri et al
EZHIP expressed
EZHIP not expressed
C
number of shared DEGs
1500
1000
500
0
1,426
396
541
1,384
up in hNPC5
down in hNPC5
up in hNPC6
down in hNPC6
3,503
3,691
4,120
4,143
0
2000
4000
total number of DEGs
D
epithelial mesenchymal transition
enrichment score
0.0
-0.5
ES = -0.54
FDR q = 0.000
E
biological process
neuron cell adhesion
Ca-dependent cell-cell adhesion via plasma membr adhes molecules
synaptic membrane adhesion
myoblast fusion
positive regulation of excitatory postsynaptic potential
negative chemotaxis
macrophage differentiation
homophilic cell adhesion via plasma membrane adhesion molecules
synapse assembly
collagen fibril organization
GO:0007158
GO:0016339
GO:0099560
GO:0007520
GO:2000463
GO:0050919
GO:0030225
GO:0007156
GO:0007416
GO:0030199
0.0
0.5
1.0
1.0
odds ratio
p-value
1e-6
1e-9
1e-12
F
DAPI
NLGN3
NLGN3
DAPI

**Figure S4. EZHIP expression is associated with neuronal-like gene programs.**
**(A-B)** Analysis of scRNA-seq data previously published by Braun et al (A) and Bhaduri et al (B) indicates that *EZHIP* is expressed in exceedingly rare cells in the prenatal human brain.
**(C)** Summary of differential gene expression analysis between EZHIP-expressing hNPCs and controls. Two hNPC cultures (hNPC5 and hNPC6) were used for these experiments.
**(D)** GSEA analysis identified negative enrichment of the "epithelial mesenchymal transition" gene set in EZHIP over-expressing hNPCs.
**(E)** GO analysis of biological processes enriched in differentially expressed genes following *EZHIP* expression.
**(F)** Control immunofluorescence experiment with no primary antibody. No NLGN3 immunofluorescence was detected. Scale bar: 20 μm.
